# Dual strategies in human confidence judgments

**DOI:** 10.1101/2020.09.17.299743

**Authors:** Andrea Bertana, Andrey Chetverikov, Ruben S. van Bergen, Sam Ling, Janneke F.M. Jehee

## Abstract

Although confidence is commonly believed to be an essential element in decision making, it remains unclear what gives rise to one’s sense of confidence. Recent Bayesian theories propose that confidence is computed, in part, from the degree of uncertainty in sensory evidence. Alternatively, observers can use physical properties of the stimulus as a heuristic to confidence. In the current study, we developed ideal observer models for either hypothesis and compared their predictions against human data obtained from psychophysical experiments. Participants reported the orientation of a stimulus, and their confidence in this estimate, under varying levels of internal and external noise. As predicted by the Bayesian model, we found a consistent link between confidence and behavioral variability for a given stimulus orientation. Confidence was higher when orientation estimates were more precise, for both internal and external sources of noise. However, we observed the inverse pattern when comparing between stimulus orientations: although observers gave more precise orientation estimates for cardinal orientations (a phenomenon known as the oblique effect), they were more confident about oblique orientations. We show that these results are well explained by a strategy to confidence that is based on the perceived amount of noise in the stimulus. Altogether, our results suggest that confidence is not always computed from the degree of uncertainty in one’s perceptual evidence, but can instead be based on visual cues that function as simple heuristics to confidence.

## Introduction

Confidence is an essential element of decision-making. When deciding whether to cross a busy intersection, for instance, we wait until confident that we will make it safely to the other side. We do not wave to a person in the distance until we are sure enough that they are our friend. And we only eat a fruit if confident that it has not gone bad. What computations give rise to this sense of confidence? Recent Bayesian theories propose that confidence is computed, in part, from the degree of uncertainty in sensory information (Aitchison, Bang, Bahrami, & Latham, 2015; Hangya, Sanders, & Kepecs, 2016; Kepecs & Mainen, 2012; Meyniel, Sigman, & Mainen, 2015; Pouget, Drugowitsch, & Kepecs, 2016). More specifically, these theories propose that confidence is a function of the posterior probability of being correct, which links confidence directly to the reliability of the evidence on which a decision is based. Thus, greater imprecision (uncertainty) in sensory evidence reduces the probability that a perceptual choice is correct, which should result in lower levels of confidence. Some support for this notion has recently been found (Barthelme & Mamassian, 2009; Navajas et al., 2017; Sanders, Hangya, & Kepecs, 2016). However, there is also an alternative interpretation. Most previous studies on confidence have varied the level of external noise in the stimulus to manipulate the reliability of sensory evidence. This is problematic, because it could be that observers simply monitor these stimulus properties as external cues to confidence (Barthelme & Mamassian, 2010; van Bergen, Ma, Pratte, & Jehee, 2015). A low contrast stimulus, for instance, typically results in poorer perceptual performance, so contrast could serve as a reliable cue to the correctness of the decision. Indeed, some physical image properties, such as variability across multiple elements of the stimulus, have been shown to impact confidence judgments (Boldt, de Gardelle, & Yeung, 2017; de Gardelle & Mamassian, 2015; Spence, Dux, & Arnold, 2016; Zylberberg, Roelfsema, & Sigman, 2014).

In the current study, we ask whether confidence is consistent with Bayesian computations involving sensory uncertainty, or rather reflects such heuristic strategies. To address this question, we capitalize on a well-studied ‘oblique effect’ in perception: orientation judgments tend to be more precise for cardinal (horizontal and vertical) than oblique orientation stimuli (Appelle, 1972; Girshick, Landy, & Simoncelli, 2011; Tomassini, Morgan, & Solomon, 2010). This phenomenon is particularly well suited to the issue at hand because it enables us to create a paradoxical situation, in which physical stimulus noise (which can be used as a cue to confidence) is more discriminable at cardinal orientations, whereas sensory uncertainty is greater for oblique orientations. This situation, as we will illustrate in simulations below, enables us to disentangle the role of heuristics from Bayesian computations in perceptual decision-making.

We will first present the theoretically ideal observer who computes confidence from the degree of uncertainty in perceptual information. The model predicts that greater levels of confidence should be linked to less variable behavior, as behavioral variability is linked to evidence reliability. We will then contrast this Bayesian model with one that uses a heuristic strategy to estimate confidence based on perceived amounts of noise, and test both models in behavioral experiments. In the experiments, participants viewed a noisy orientation stimulus that consisted of multiple elements. The orientation of the elements was varied to manipulate noise. Participants reported both the perceived (global) orientation across the elements, and their confidence in this estimate (Figure 1). Consistent with the Bayesian model, we found that for a given stimulus orientation, confidence reliably tracked behavioral performance. Specifically, orientation judgments were more precise with higher confidence, for fluctuations in both external and internal noise. Surprisingly, however, the results deviated from Bayesian predictions when comparing between stimulus orientations: although orientation estimates were more precise for cardinal (horizontal and vertical) than oblique orientations (Appelle, 1972; Girshick et al., 2011; Tomassini et al., 2010), confidence was *higher* for oblique orientations. Rather than being consistent with Bayesian decision theory, we show that these results are better explained by the participant’s ability to perceive the noise in the stimulus – a heuristic to confidence.

**Figure 1.**
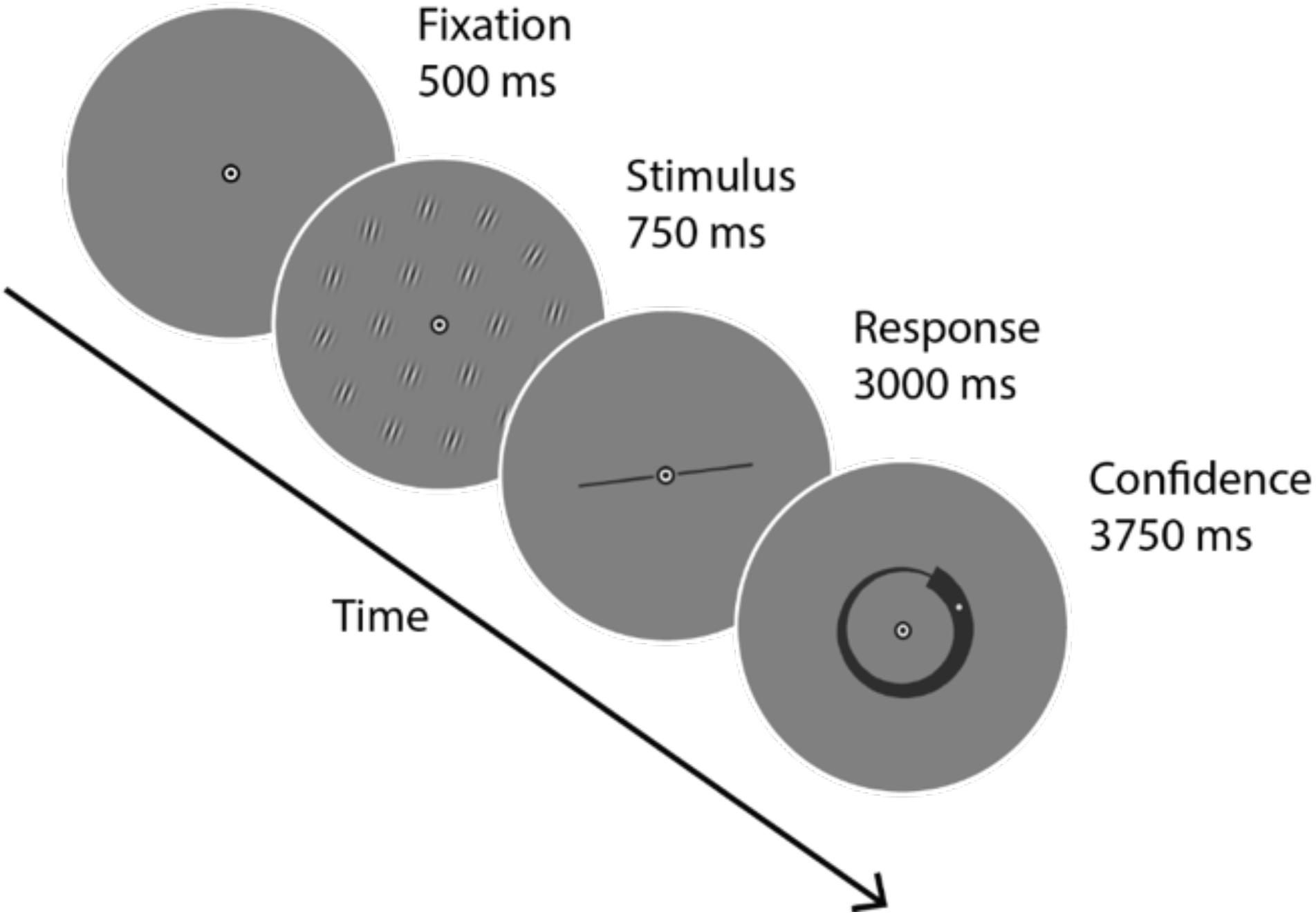
Schematic representation of a single trial in Experiments 1 and 2. The number, size and position of the elements in the stimulus, as well as the size of the bar and spiral are not an exact copy, but were adapted for illustrative purposes. On each trial, subjects first reported the mean orientation across 36 Gabor patches by rotating a bar that appeared around fixation. They then reported their confidence in this orientation estimate by rotating a dot within a spiral. In Experiment 1, sensory uncertainty (orientation variability) was manipulated using five noise levels. In Experiment 2, a single level of orientation noise was used and the mean orientation of the stimulus was manipulated.

## Methods

### Participants

10 healthy adult volunteers (aged 18-35, six female) participated in Experiment 1, and 8 healthy adult volunteers participated in Experiments 2 (aged 19-35, five female) and 3 (aged 22-35, five female). Seven of the observers of Experiment 1 also participated in both experiments 2 & 3. All participants had normal or corrected-to-normal vision, and all gave informed written consent. The study was approved by the Radboud University and Boston University Institutional Review Boards.

### Apparatus

The same set-up was used across two experimental sites. The stimuli were generated by a Macbook Pro running Matlab 2014b and the Psychtoolbox software package (Brainard, 1997; Pelli, 1997), and displayed on a gamma-calibrated CRT monitor (resolution: 1024 x 768 pixels; refresh rate: 120 Hz). A black annulus was positioned around the screen to prevent subjects from using the edge of the frame as a reference orientation in the task.

### Experimental design

#### Experiment 1 – five levels of external noise

Participants were instructed to maintain fixation at a central bullseye target (radius: 0.5° of visual angle (v.a.)) throughout the duration of the experiment. A schematic representation of a single trial is shown in Figure 1. Each trial started with an initial fixation period (duration: 500 ms). This was followed by the stimulus (duration: 750 ms), after which participants gave their response (duration: 3000 ms). Subjects reported the mean orientation of the elements of the stimulus by rotating a bar (radius: 2.5°) presented at fixation. The bar was presented at an initially random orientation, and participants adjusted its orientation using a mouse. Their responses were recorded when they clicked the left mouse button. Directly after reporting the orientation of the stimulus, participants indicated their confidence in their orientation judgment (duration: 3750 ms) by positioning a small dot on a black spiral presented around fixation. The width of the spiral varied linearly and indicated the degree of confidence. The spiral appeared at an initially random orientation and direction (clockwise vs. counterclockwise), and the dot’s initial orientation within the spiral was determined randomly. We counterbalanced the direction of the confidence scale across subjects, such that for some subjects, high confidence corresponded to the region where the spiral was broader, while for others it corresponded to the region where the spiral was narrower. Confidence reports had a range of 0 (minimum confidence) to 1 (maximum confidence). The participants adjusted the dot’s location using a mouse, and their responses were recorded when they clicked the left mouse button.

The stimulus consisted of a pattern of 36 full-contrast Gabor patches, presented on a uniform gray background. Each Gabor element was comprised of a sinusoidal grating (4 cycles/°) masked by a circular Gaussian (full width at half maximum: 0.5° of v.a.). The orientation *θ*_*tj*_ of the *j*-th Gabor element on the *t*-th trial was determined as follows:

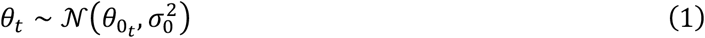

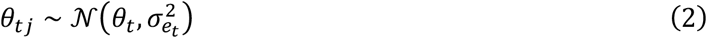

where 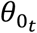 refers to the base orientation that was (randomly) selected for trial *t* from one of four possibilities (22.5°, 67.5°, 112.5° or 157.5°), *θ*_*t*_ is the mean stimulus orientation on trial 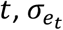 is the external noise level on this trial (standard deviation of Gaussian noise, chosen from 0.5°, 2°, 4°, 8°, or 16°) and σ_0_ = 7° models a small jitter of the base orientation across trials. In words, we first created 20 conditions, one for each combination of base orientation (22.5°, 67.5°, 112.5°, 157.5°) and noise level (standard deviation of Gaussian noise: 0.5°, 2°, 4°, 8°, 16°). For each stimulus condition, we then created 105 orientation trials by adding small orientation offsets to the condition’s base orientation (the offsets are randomly sampled from a Normal distribution, 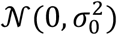) to reduce predictability. We then generated, for each trial, 36 patches using a Gaussian distribution centered on that trial’s stimulus orientation, and with noise level determined by the stimulus condition. Each of these Gabor patches was subsequently positioned on a grid defined by three concentric rings (of radius 2°, 4° and 6° v.a.) around the fixation target (the grid was not visible to participants). Individual elements were first positioned evenly within each ring in the grid, and then random jitter (uniformly distributed; min = -0.75°, max = 0.75°) was added to both polar coordinates of the grating’s location, such that spatial predictability was reduced.

Subjects completed 60 blocks of 35 trials each for a total of 2100 trials per subject. Within each block, we used one level of orientation noise. Each subject completed one experimental session per day. For two subjects there were 3 sessions of 20 blocks each, while for the remaining eight subjects there were 6 sessions of 10 blocks each. In each session, all five levels of orientation noise were shown at least twice in random order. Prior to the start of the experiment, each observer practiced the task on a separate day by completing at least one block of each level of orientation noise used in the experiment.

#### Experiment 2 – all orientations

Experiment 2 used the exact same procedure and stimuli as Experiment 1, except that the mean (base) orientation of the stimulus was pseudo-randomly sampled from 0-180°, and we used only a single noise level on individual stimulus elements and no offset on the base orientation. The stimulus consisted of 36 Gabor patches that were drawn from a Normal distribution with mean equal to the (base) orientation of that trial and standard deviation set to 0.5°. For every experimental block, base orientations were spaced every 4 degrees to ensure uniform coverage of the orientation space. The starting orientation for this selection was randomly picked out of four alternatives ([1°, 2°, 3°, 4°]), so that across blocks, stimuli covered the orientation space every 1°. This resulted in 46 trials for each block, in which orientations were presented in random order. Each subject completed 16 blocks of 46 trials each, for a total of 736 trials divided over two days. One observer did not participate in Experiment 1; this observer practiced the task for 4 experimental blocks prior to the start of the experiment.

#### Experiment 3 – variance discrimination

Participants were instructed to maintain fixation at the central bullseye target (radius = 0.5° of v.a.) throughout the experiment. To indicate the start of each trial, the fixation bullseye briefly turned black (duration: 500 ms), after which the first stimulus appeared on the screen (duration: 750 ms), followed by an interval (duration: 500 ms), and then a second stimulus (duration: 750 ms). After the second stimulus disappeared from the screen, subjects indicated which of the two stimuli contained more orientation noise by pressing the ‘1’ or ‘2’ button on a keyboard (duration: 1000 ms). As in Experiments 1 & 2, the orientation stimuli consisted of 36 Gabor patches. The orientation of the patches was drawn from a Gaussian distribution centered on one of twelve base orientations (0-180°, every 15°). To ensure the trial mean was always identical to the base orientation, we first created 10,000 stimuli by randomly sampling from the Gaussian distribution, and then selected those stimuli for which the sample mean across the Gabor patches approximated the true mean with error smaller than 0.001°. For one of the two stimuli presented on each trial, the standard deviation of the Gaussian distribution was always 0.5°. For the second stimulus, the standard deviation of the Gaussian distribution was determined using a staircase procedure (2 down – 1 up, starting at 4° standard deviation). The two stimuli always had the same mean orientation, and their presentation order was randomized across trials. Patch location was determined independently for the two stimuli, and using identical procedures as in Experiments 1 & 2. After 2 experimental blocks of practice, participants completed 36 blocks of 50 trials each, over 3 days. On each day, all twelve orientations were presented in random order; however, within one block only one mean orientation was shown.

### Data analysis

#### Experiments 1 & 2

For both experiments, behavioral error was computed as the acute-angle distance between the reported orientation and the mean of the generative distribution of the stimulus on that trial. Behavioral variability was calculated as the across-trial variance of the behavioral errors. In Experiment 2, behavioral variability was calculated after correcting for an orientation-dependent shift in mean (i.e., a bias away from the cardinals; see also (van Bergen et al., 2015) by fitting a 4-degree polynomial to the behavioral errors (note that in Experiment 1, the presented orientations were at roughly the same distance to the cardinals, 22.5°, so no such correction was necessary). We used the residuals of this fit in our calculations of behavioral variability. Trials for which the behavioral error of the participant exceeded the mean error by more than +/-3 times the standard deviation across trials (computed for each subject separately) were excluded from further analysis, as this large an error suggests that the observer randomly guessed the orientation of the stimulus. This occurred on 0.81% of all trials in Experiment 1, and 0.51% of all trials in Experiment 2. Control analyses verified that our conclusions remain the same when including all trials for further analysis. Despite our instruction that participants use the whole range of the confidence scale available, some observers gave confidence ratings that were mostly restricted to a smaller range. To account for this variability in the distribution of confidence reports across subjects, confidence reports were z-scored for each subject within each experiment.

To investigate the impact of external noise on behavior, trials from Experiment 1 were sorted, for each observer, into five equal-sized bins based on the amount of orientation noise in the stimulus. Behavioral variability (standard deviation; *SD*_beh_) and mean level of confidence across all trials in each of the bins was computed in two steps: first, for each (base) orientation separately, and then averaged over orientations. To test for significance, a linear regression was used:

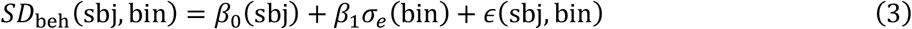

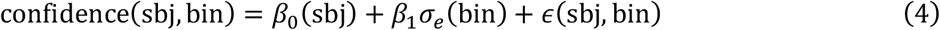

where *β*_0_(sbj) is a subject-specific intercept term, σ_*e*_(bin) is the amount of external stimulus noise in a given bin, *β*_1_ is the associated regression coefficient, and *ϵ*(sbj, bin) is the residual error for a given subject × bin (resulting in a total of 11 independent variables for each model: 10 subject-specific intercepts and 1 slope coefficient on the external noise). *β, β*_0_(sbj), and *ϵ* were estimated separately for confidence and behavioral variability.

To assess the impact of internal noise on behavioral variability and reported confidence in Experiment 1, the effect of mean stimulus orientation, as well as local changes in variance across the 36 stimulus elements (i.e., small trial-by-trial changes in the sample variance), on reported confidence were first removed via linear regression for each observer:

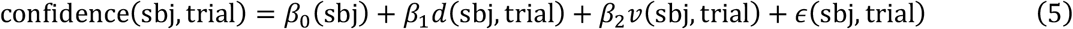

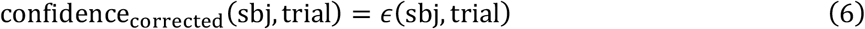

where *β*_0_(sbj) is a subject-specific intercept, *d* is the mean stimulus orientation expressed as distance to the nearest cardinal (i.e., horizontal or vertical) for each subject on each trial, *v* is the variance across the 36 patches, *β*_1_ and *β*_2_ are the corresponding regression coefficients, and *ϵ*(sbj, trial) is the residual error. The residuals of this analysis, that is, confidence after removing the effects of mean stimulus orientation and local changes in variance across patches (confidence_corrected_ = *ϵ*(sbj, trial)), were then used to sort trials into three bins of increasing confidence for each observer and each of the five external noise levels independently. Behavioral variability was computed for each of these bins, resulting in three values of behavioral variability (one for each level of confidence) for each level of external noise. To test for significance, a two-way ANOVA was conducted on behavioral variability values, with external noise and confidence as factors. This was followed by a sequential comparison between the low and medium and medium and high confidence levels, using an ANOVA with the same two factors. To test whether confidence predicted behavioral variability when external noise was low, behavioral variability was linearly regressed on confidence, using data for the lowest external noise level only:

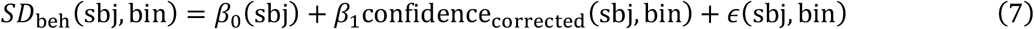

where *β*_0_(sbj) is a subject-specific intercept term, confidence_corrected_(sbj, bin) is the average confidence within each confidence bin (corrected as explained above), *β*_1_ is the associated regression coefficient, and *ϵ*(sbj, bin) is the residual error for a given subject × bin (resulting in a total of 11 independent variables for each model: 10 subject-specific intercepts and 1 slope coefficient for confidence).

To test the impact of internal noise on orientation perception and confidence in Experiment 2, data were sorted into 12 equally spaced orientations bins, each covering a 15° range with the first bin centered at 0°. Behavioral variability was computed across all trials in each orientation bin. To test whether behavioral variability, or reported levels of confidence, reliably changed across orientation, a linear regression was used:

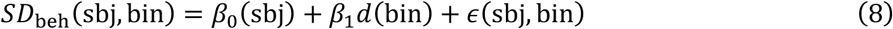

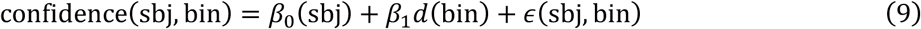

where *d*(bin) is the orientation of the stimulus within a bin, defined in terms of its distance to the nearest cardinal orientation (i.e., horizontal or vertical). The regression models also included an intercept for each individual observer, *β*_0_(sbj), resulting in a total of 9 independent variables in each model.

#### Experiment 3

We estimated the amount of internal noise across orientation stimuli using the approach of Morgan et al. (Morgan, Chubb, & Solomon, 2008), which is based on signal detection theory. Subjects were presented with pairs of stimuli separated by a fixed interval, and were asked to report which stimulus had the higher variance across *n* = 36 stimulus elements (2-alternative forced choice). The elements of the first stimulus (standard) were drawn from a Gaussian distribution with variance 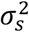, while those of the second (comparison) stimulus were drawn from a Gaussian distribution with variance (σ_*s*_ + Δσ)^2^. Because the elements were drawn from a Gaussian distribution, the variance across the *n* stimulus elements followed a scaled χ^2^ distribution across trials, with *n*-1 degrees of freedom. Adding internal noise 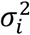, the probability of reporting that perceived variance of the standard 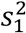 is greater than that of the comparison 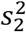 becomes:

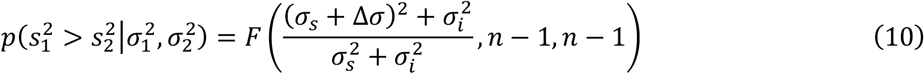

where *F* is the cumulative distribution function of the *F*-distribution, with degrees of freedom *n*-1 and *n*-1. See (Morgan et al., 2008) for more detailed derivations. We estimated internal noise value 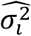 for each participant and base orientation independently, using maximum-likelihood estimation (MLE) and the data of Experiment 3.

### Observer models

#### Orientation judgments (both models)

We start with the observation that sensory signals are corrupted by noise. Thus, even when the external stimulus is held constant, sensory signals vary from trial to trial. For a given stimulus, this results in a distribution of possible signals, or measurements, which is called the measurement distribution *p*(*m*|*θ*). We assumed that the measurements are drawn from a Normal distribution centered on *θ*, with variance determined by both internal and external sources of noise, 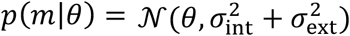. For the compound stimulus considered here, both sources of noise are directly related to the across-trial variance in individual stimulus elements (see *Simulations*). External noise refers to noise in the environment, and we considered two sources of noise internal to the observer. The first source of internal noise was independent of stimulus orientation, and could be high, medium or low. The second source varied across orientation stimuli, with smaller levels of noise for cardinal (horizontal and vertical) than oblique orientations. This pattern of internal noise models an oblique effect in orientation perception (Appelle, 1972; Girshick et al., 2011; Tomassini et al., 2010).

The observer’s task is to infer the orientation of the stimulus from incoming sensory signals. The ideal observer uses knowledge of the measurement distribution, combined with Bayes’ rule, to infer the range of stimuli *p*(*θ*|*m*) that could have led to the current measurement. Assuming a flat stimulus prior, this becomes:

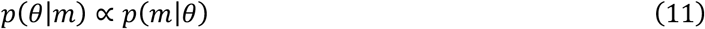

This posterior distribution represents the observer’s beliefs about which stimulus orientations are likely given the current measurement. We take the mean of the posterior distribution as the model observer’s estimate of orientation. The variance of the posterior distribution 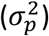 can be taken as a measure of the degree of uncertainty in this estimate. Additional simulations verified that if observers learn the task-specific distribution over orientation (which was non-uniform in Experiment 1) and use it as a prior instead of a flat prior as assumed here, the model predictions are qualitatively the same.

#### Bayesian model of confidence

The Bayesian confidence hypothesis holds that confidence is based on the degree of sensory uncertainty. More specifically, this hypothesis predicts that confidence should be inversely related to the degree of sensory uncertainty. In our simulations, we quantified this inverse relationship between confidence and uncertainty as follows:

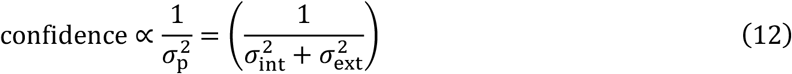

Note that this equation also describes the amount of Fisher information contained in the observations. See Navajas et. al. (2017) for a similar definition of confidence.

#### Heuristics model of confidence

The Heuristics model observer uses the same decision-making process as the Bayesian model to report the orientation of the stimulus. However, rather than basing confidence on uncertainty, this model assumes that observers compute confidence from the perceived amount of external noise in the stimulus. Because the stimuli consist of arrays of Gabor functions (see Fig. 1), we define perceived noise as the variance across the physical stimulus elements, as estimated from the observer’s measurements on a given trial.

Recall that the noisy stimulus elements are drawn from a Normal distribution around stimulus orientation *θ*, and with variance 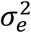. We assume that on each trial, the observer takes a set of orientation measurements {*m*_*j*_} from this compound stimulus. When the observer takes the sensory measurements, internal noise adds further variability. Thus, the sensory measurement *m*_*j*_, of the orientation of the *j*-th stimulus patch, follows a Normal distribution:

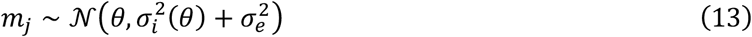

where 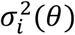 is the internal noise variance in the measurements of individual orientation patches, as a function of the overall stimulus orientation *θ*.

We assume that the observer has learned, from prior experience, how internal noise varies as a function of stimulus orientation; that is, the observer knows 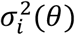. Note that we do not assume that the observer knows the actual internal noise level at a given trial, just the across-trial estimate of the level of internal noise for a given orientation. Given this information, and combined with the set of orientation measurements {*m*_*j*_}, the observer computes an estimate of the amount of external noise in the stimulus. The observer does this by maximum-likelihood estimation (MLE), finding the value of σ_*e*_ that maximizes the probability of the observed data (the measurements {*m*_*j*_}):

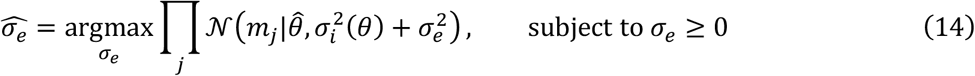

where 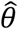 is the observer’s estimate of the stimulus orientation on that trial, computed by averaging the measurements of individual elements in the set (i.e., 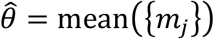). Given the Normal distribution of the measurements, and the constraint that σ_*e*_ must be strictly non-negative (or in other words, that the sample variance cannot be exceeded), the following equation to compute 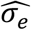 can be derived analytically:

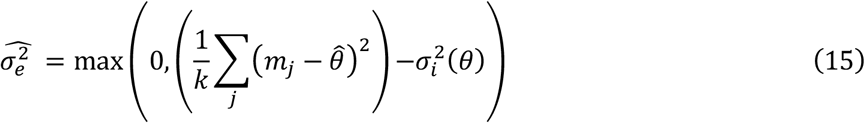

where *k* is the number of measurements taken by the observer, which may be fewer than the total number of stimulus elements *n*. In words, the observer computes the sample variance of the orientation measurements {*m*_*j*_}, and subtracts the variance that can be explained by internal noise, to obtain a perceptual estimate of external noise variance. If the entire sample variance can be accounted for by internal noise, then the perceived external noise variance is 0.

The Heuristics observer uses this perceived external noise variance to compute confidence:

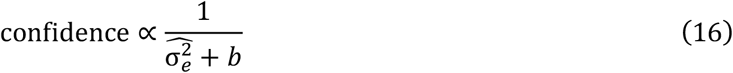

where *b* is a parameter that determines the observer’s maximum confidence (when 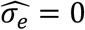). Thus, the Heuristics observer’s confidence is inversely proportional to the estimated external noise variance: the more noise the observer perceives in the stimulus, the less confident the observer will be. Note that this model is not normative, so we simply defined confidence as a function of the amount of perceived noise in the stimulus. While the estimated amount of stimulus noise tends to be more accurate when the observer takes more measurements from the stimulus, this perceived noisiness (and hence confidence) will not decrease (increase) with larger *k*. However, had we nonetheless incorporated the *k* parameter into Equation 16 (thereby creating higher levels of confidence when the observer samples more orientation patches), this would have merely scaled the predictions, and their overall pattern would have remained the same.

### Simulations

We simulated the behavioral estimates of the two model observers in two experiments. The simulations of Experiment 1 used a single orientation stimulus *θ*, while for Experiment 2 we randomly drew orientations from a uniform orientation distribution (from 0 to 180°). We assumed that the across-trial variance 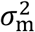^2^ in the observer’s measurements *m* arose due to sources both internal and external to the observer (see also above):

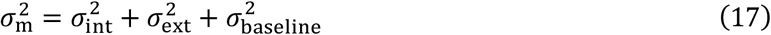

where 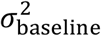 is a baseline parameter to account for downstream sources of noise. We assume that the observer optimally integrates the measurements of *k* individual stimulus elements to calculate the most likely mean stimulus value, so that behavioral variability due to external noise alone is given by 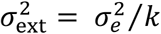. In line with the actual experimental parameters (see *Experimental Design*), we set σ_*e*_ to 120 linearly spaced values from 0.5° to 16° for our simulations of Experiment 1 (Figure 2a, b, e, f), and to 0.5° for Experiment 2 (Figure 4a, b). Similarly, we assumed that 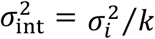, with internal noise σ_*i*_ set to 6°, or either low, medium or high (i.e., 4°, 6° and 8°) for our simulations of Experiment 1 (Figure 2a & b, and 2e & f, respectively). We used 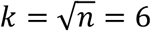 in our simulations, as human observers typically pool across approximately the square root of the *n* stimulus elements for their decisions (Dakin, 2001; Moerel, Ling, & Jehee, 2016; Whitney & Yamanashi Leib, 2018). However, using a different value of *k* does not qualitatively change any of our predictions (Figure S3). For the simulations of Experiment 2 (Figure 4a, b), internal noise was dependent on stimulus orientation, with smaller levels of noise for cardinal than oblique orientations:

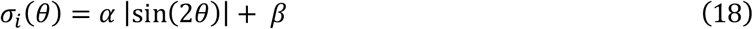

where *θ* is the presented orientation in radians (where horizontal is *θ* = 0 rad), *α* = 7 is an amplitude parameter, and *β* = 1 implements the baseline at cardinal orientations. This pattern of internal noise models the well-known oblique effect in orientation perception (Appelle, 1972; Girshick et al., 2011; Tomassini et al., 2010). To illustrate the relationship between internal noise and confidence and behavioral variability (Fig. 2c, d), σ_*i*_ was set to 120 linearly spaced values between 0.5° and 16°, and σ_*e*_ was 6°. For all simulations, we added a constant baseline of 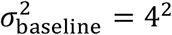 (Equation 17); note that this parameter does not affect the overall pattern of effects in the simulated data. For both model observers, the reported stimulus orientation (i.e. their behavioral estimate) was obtained using Equation 11 and taking the (circular) mean of the distribution, as outlined above. Confidence estimates were obtained using Equations 12 and 16, for the Bayesian and Heuristics model observer, respectively. For each model observer, confidence values were normalized across experiments to lie in between 0 (minimum confidence) and 1 (maximum confidence).

**Figure 2.**
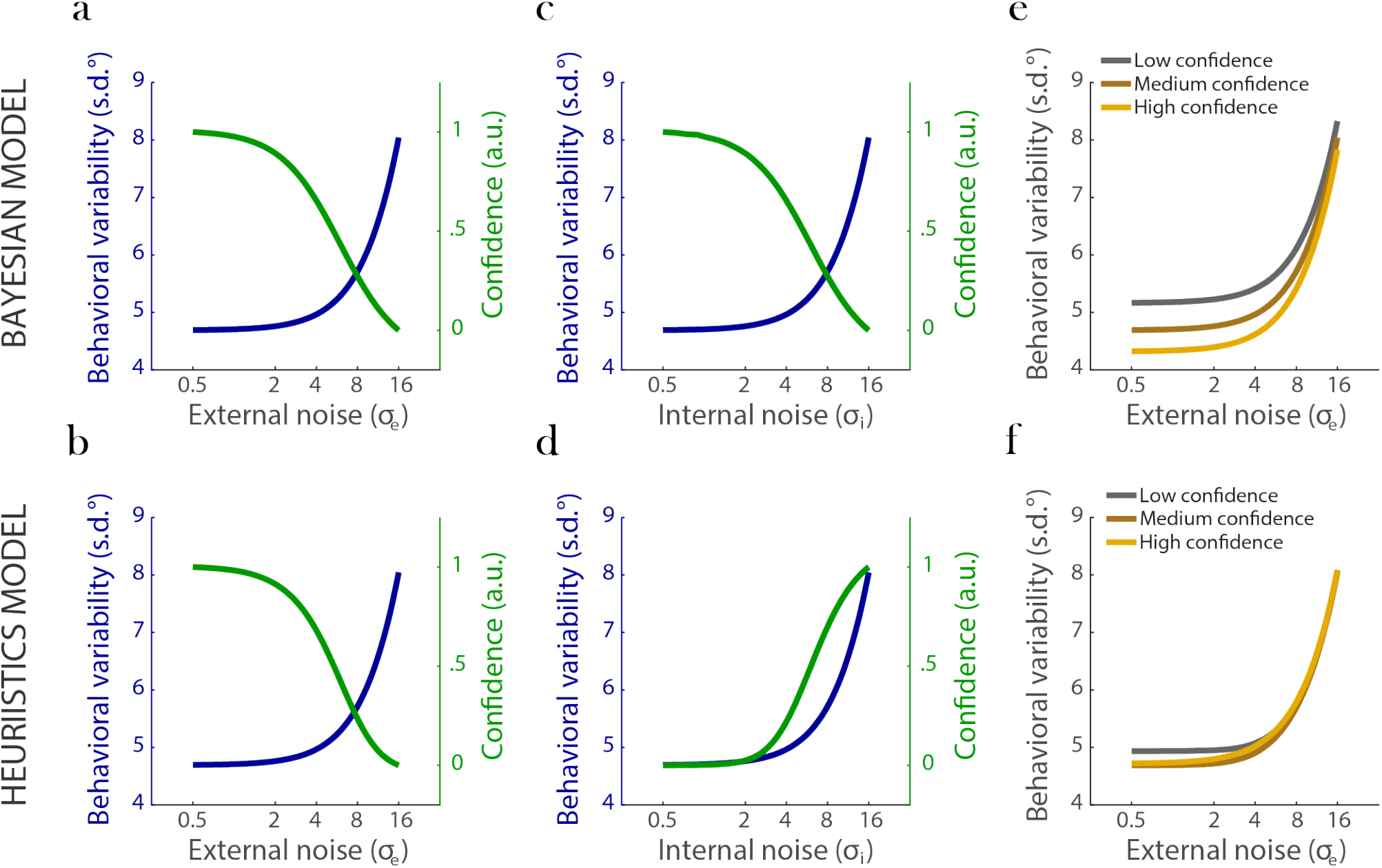
Predictions for the Bayesian and Heuristics model observers. (**a-d**) Variability in behavioral orientation estimates and mean reported confidence as a function of external (**a-b**) and internal (**c-d**) noise, for the Bayesian (**a, c**) and Heuristics (**b, d**) model observer. For both observers, behavioral variability increases with increasing levels of external and internal noise. In addition, confidence decreases with increasing levels of external noise. Thus, either model observer predicts lower levels of confidence when behavioral performance worsens due to external sources of noise. The predictions of the two models diverge when internal sources of noise are considered. The Bayesian observer reports lower confidence when internal noise increases, such that confidence and behavioral variability remain inversely linked. In contrast, the Heuristics observer paradoxically becomes *more* confident when internal noise increases, even though behavioral performance deteriorates. (**e-f**) Predictions when trials are sorted with respect to reported confidence. For each level of external noise, trials were sorted into three bins of increasing confidence, and behavioral variability was computed across all trials in each bin. For the Bayesian observer (**e**), confidence correctly predicts behavioral variability: when the observer reports high confidence, orientation estimates tend to be less variable. For the Heuristics observer (**f**), the relationship between confidence and behavioral performance is less straightforward, as the effects of external and internal noise compete. In all figures, the x-axis is plotted using a logarithmic scale.

**Figure 3.**
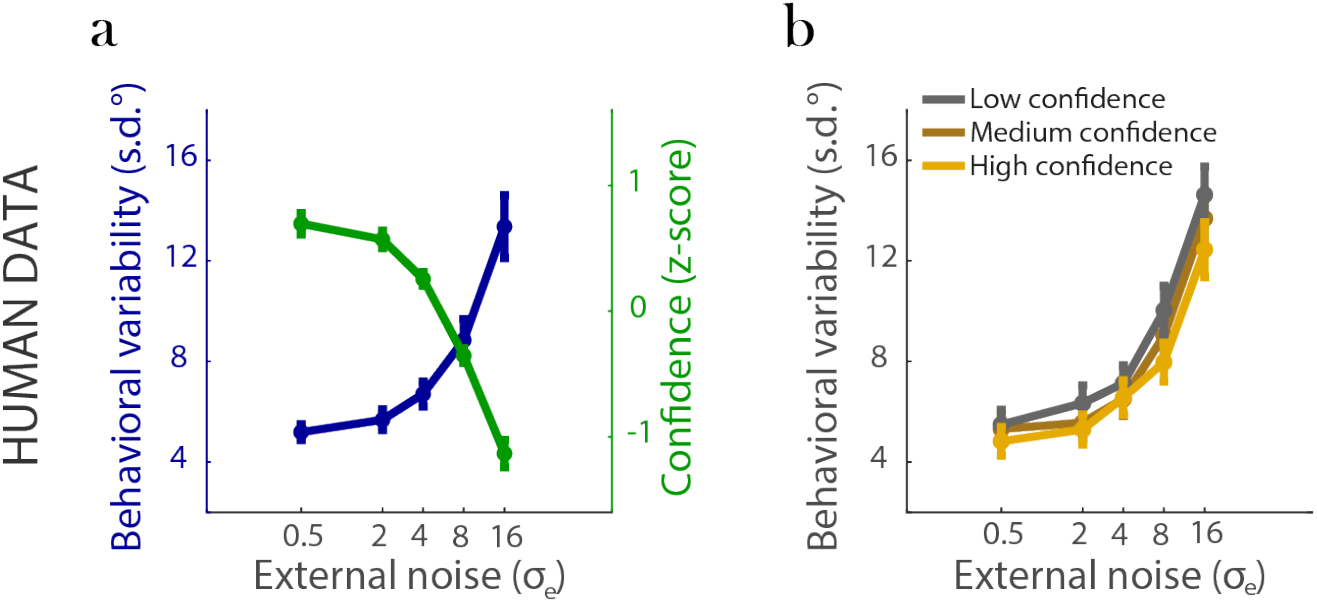
Variability in human behavioral orientation estimates, and associated levels confidence, across various levels of external noise. (**a**) Behavioral variability (blue) increases with increasing levels of orientation noise in the stimulus, while reported confidence (green) decreases. (**b**) For each level of external noise, the data were divided into 3 bins of increasing confidence (black, brown and yellow lines), and behavioral variability was computed across all the trials in each bin. Data were averaged over (base) orientations. As can be seen in the figure, behavioral variability decreased when reported confidence increased (the yellow line generally lies below the brown and black lines). Error bars represent ±1 s.e.m.

**Figure 4.**
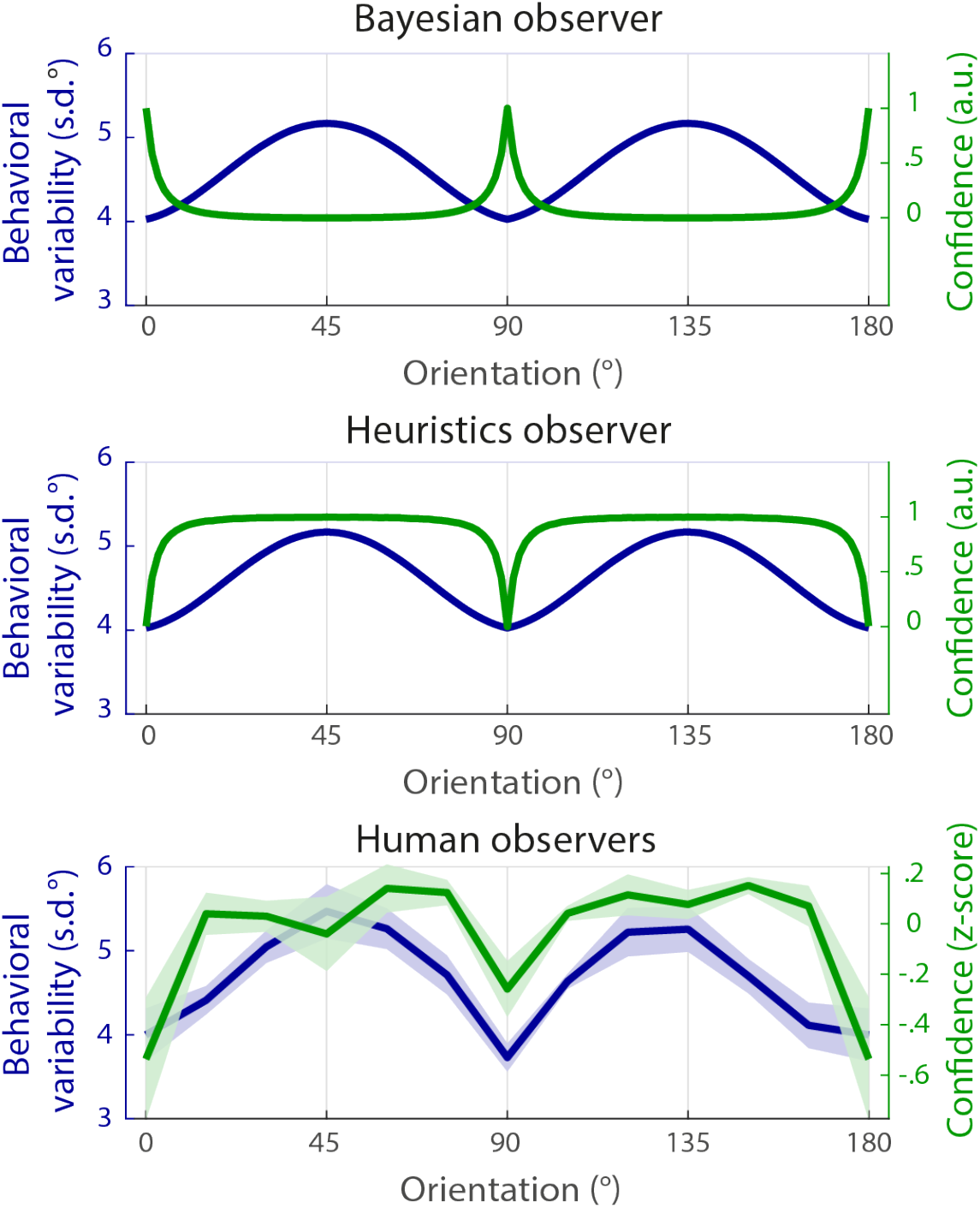
Behavioral variability (blue) and reported confidence (green) across orientation stimuli (Experiment 2), for both model and human observers. **Top**: Bayesian model observer who computes confidence from the degree of uncertainty in their orientation estimates. Confidence decreases when behavioral variability increases (oblique orientations). **Middle**: Heuristics model observer who computes confidence from the amount of perceived noise in the stimulus. Confidence is positively linked to behavioral variability: when behavioral variability increases (oblique orientations), so does confidence. **Bottom**: Human observers expressed greater levels of confidence for oblique compared to cardinal orientation stimuli, whereas their orientation estimates were less precise for oblique stimuli. These results are most consistent with the predictions of the Heuristics model observer. Shaded area represents ±1 s.e.m.

To generate a quantitative prediction of confidence across levels of external noise in the cross orientation for the participants of our experiment (Fig. 5b), we first estimated internal noise values using the data of Experiment 3 (see *Data Analysis*). For each of the 12 measured orientations and for each individual observer, we then predicted their level of confidence using these estimates and Equations 12 and 16. We subsequently averaged across observers to arrive at Figure 5b.

**Figure 5.**
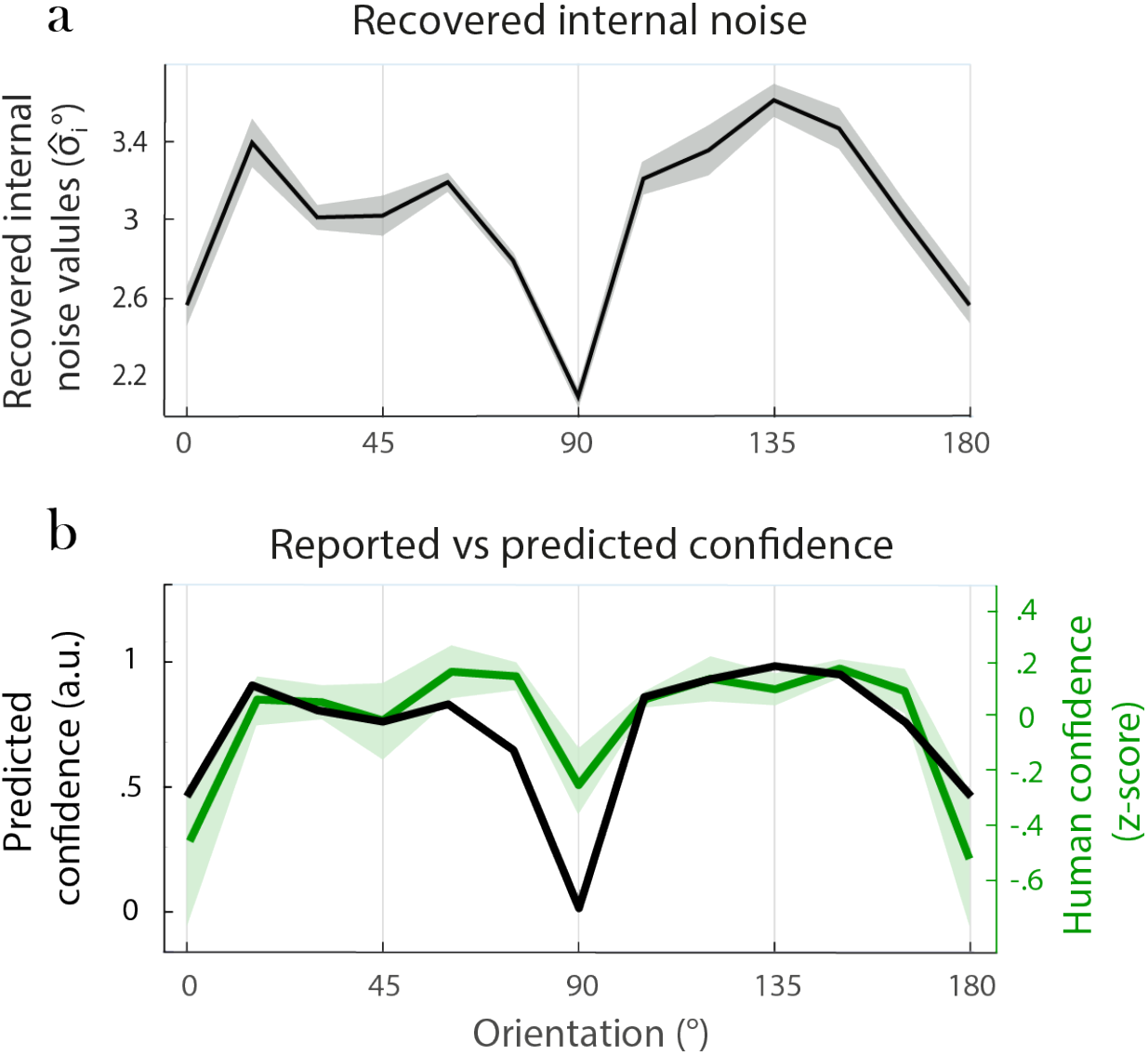
Internal noise, predicted confidence, and actual confidence. (a) Recovered internal noise values for different orientation stimuli, using Equation 10 and the data from the noise discrimination task (Experiment 3). Human observers are better able to discriminate small changes in stimulus variance for cardinal than oblique orientation stimuli, which indicates that internal noise values are higher for oblique than cardinal orientations (r = 0.48, t(87) = 5.12, p < 0.001). (b) Predicted confidence for the Heuristics model observer (black) using realistic levels of internal noise (from a), shown alongside the actual levels of confidence reported by human observers (green). Predicted confidence nicely matches reported confidence, suggesting that human observers use heuristic strategies to confidence based on the amount of perceived noise in the stimulus. Shaded area represents ±1 s.e.m.

## Results

### Confidence models

Do human confidence judgments reflect the imprecision in people’s perceptual evidence? To address this question, we first discuss the statistically ideal observer who bases confidence on the degree of uncertainty in perceptual information (Bayesian observer), followed by a model that uses a perceived-noise heuristic to confidence (Heuristics observer). We will then test several model predictions against experimental data to see which model observer best describes human behavior.

The ideal observer’s task is to infer the orientation of a stimulus from incoming sensory signals. These signals, or measurements, are corrupted by noise. Thus, for a given sensory stimulus *θ*, the observer’s measurement *m* changes from one trial to the next, and the relationship between the stimulus orientation and its measurements is described by a probability distribution, *p*(*m*|*θ*). We assume that the measurements are drawn from a Gaussian distribution centered on *θ*, with variance determined by a combination of internal and external sources of noise, 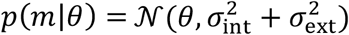 External noise refers to noise that occurs in the environment; for instance, because physical stimulus elements are variable. We model two sources of internal noise. The first source of internal noise is independent of stimulus orientation, and could be high, medium or low. It captures trial-by-trial fluctuations in, for example, the observer’s attentional state or fatigue. The second source varies with stimulus orientation, with smaller levels of noise for cardinal (horizontal and vertical) than oblique orientations. This pattern of internal noise models the well-known oblique effect in orientation perception (Appelle, 1972; Girshick et al., 2011; Tomassini et al., 2010).

When estimating the stimulus from a given measurement, the statistically ideal observer uses knowledge of the measurement distribution to infer a probability distribution over stimuli. Specifically, assuming a flat stimulus prior and using Bayes’ rule (Equation 11), we obtain the posterior probability distribution, *p*(*θ*|*m*), which describes the range of orientations that is consistent with the current measurement *m*. The mean of the posterior distribution serves as the observer’s orientation estimate, while its width (variance, 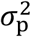) can be taken as a metric on the degree of sensory uncertainty in the estimate. The Bayesian (normative) hypothesis holds that confidence is based on the degree of sensory uncertainty. Thus, assuming that the brain computes sensory uncertainty, the question is whether or not observers are aware of this uncertainty and use it in their confidence reports. The hypothesis predicts that when the observer’s perceptual evidence is imprecise, the level of confidence should be low. In our simulations, we quantify this inverse relationship between confidence and uncertainty as follows (Equation 12):

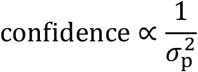

However, our predictions do not critically depend on the particular function described here, as long as confidence decreases with an increase in uncertainty (see Navajas et. al., 2017, for a similar definition of confidence).

The second model that we consider uses a heuristic strategy to confidence. Specifically, this model computes confidence from the perceived amount of external noise in the stimulus – a reasonable (albeit suboptimal) strategy, as physical stimulus noise is typically linked to the magnitude of behavioral errors. Because the stimuli consist of a set of noisy elements (see Fig. 1), we define perceived noise as the variance across these physical elements, as estimated from the observer’s measurements on a given trial.

The observer’s measurements of the stimulus elements vary from one trial to the next, and so does their variance. Specifically, the measurements *m*_*j*_ of stimulus element *j* follow a Normal distribution (see Equation 13). The variance of the distribution is described by 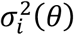, the internal noise variance in the measurements of individual orientation patches as a function of the overall stimulus orientation *θ*, and 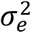, the variance in individual elements due to external sources of noise. Both sources of noise are directly related to the across-trial variance in the observer’s orientation judgements (see *Methods*). We assume that the observer has learned, from prior experience, how internal noise varies as a function of stimulus orientation; that is, the observer knows 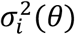. The observer uses this information and the set of measurements {*m*_*j*_} to compute, for each trial, the mostly likely estimate of the amount of external noise in the stimulus 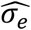 (see Equation 14). More specifically, the observer computes the sample variance of the orientation measurements, and subtracts the variance that can be explained by internal noise, to obtain the most likely estimate of external noise variance for that trial’s stimulus. If all sample variance can be accounted for by internal noise, then the perceived external noise variance is 0 – in other words, the stimulus ‘looks’ like it contained no external noise.

The Heuristics observer uses this perceived amount of external noise to compute confidence (Equation 16):

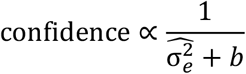

where *b* is a parameter that determines the observer’s maximum confidence (when 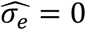). Thus, the Heuristics observer’s confidence is inversely proportional to the estimated external noise variance: the more noise the observer perceives in the stimulus, the less confident the observer will be. See *Methods* for more detailed definitions and derivations. See Supporting Materials for a third observer model, which computes confidence from the ability to *detect* noise, rather than the perceived amount of noise, in the stimulus. Its predictions are qualitatively similar to the ones obtained from the perceived-noise model observer discussed here.

### Predictions

What pattern of results should we expect for the two strategies to confidence? To illustrate the relationship between sensory uncertainty, behavioral variability and confidence, we parametrically varied internal and external sources of noise, and simulated both behavioral orientation estimates and levels of confidence for each model observer.

We first focus on behavioral orientation estimates. We computed the across-trial variability in behavioral orientation estimates across varying levels of sensory uncertainty. For either observer, behavioral variability increases with increasing levels of uncertainty. This relationship holds no matter the source of noise that creates sensory uncertainty. Thus, behavioral variability not only increases with increasing levels of external noise in the stimulus (Figure 2a-b), but also when internal levels of noise are larger, either for a single orientation (Figure 2c-d) or between orientations, as for cardinal versus oblique orientation stimuli (Figure 4 top).

What level of confidence does each model observer predict across levels of internal and external noise? Because the Bayesian observer computes confidence from the degree of uncertainty in the data, confidence is linked to the variability in the observer’s behavioral orientation estimates. This relationship holds regardless of the source of noise that we consider. Thus, for this model observer, confidence is not only (inversely) linked to behavioral performance for external sources of noise (Figure 2a), but also for internal sources of variance, both within (Figure 2c) and across orientations (Figure 4a). For the Heuristics observer, the predictions are similar to the Bayesian observer when considering levels of external noise. That is, for external noise, this model observer simply perceives the change in physical stimulus properties, and adjusts confidence levels accordingly. Much like the Bayesian observer, this results in decreasing levels of confidence with greater levels of stimulus noise or uncertainty (Figure 2b). Interestingly, however, the predictions deviate between the two observers across different levels of internal noise. Because an increase in internal noise hampers the ability of the Heuristics observer to accurately detect variability in the physical stimulus elements, this model observer reports high levels of confidence with decreasing behavioral performance, both within (Figure 2d) and across (Figure 4b) orientations.

In an actual (rather than simulated) experiment, the psychophysicist doesn’t have direct access to the amount of internal noise in the observer’s orientation estimates. How best to investigate the relationship between internal noise, confidence, and behavioral performance for more realistic experimental scenarios? We reasoned that when external noise in the stimulus is held constant, any change in behavior has to be caused by fluctuations in internal factors. This implies that if confidence is somehow based on these internal factors, it should be linked to behavioral performance, as well. Following this logic, we sorted trials in bins of increasing confidence and computed behavioral variability for each of the bins. What pattern of results should one expect for such an analysis within orientation stimuli? For the Bayesian model observer, reported levels of confidence again predict the degree of precision in orientation estimates, both within and across levels of external noise (Figure 2e). For the Heuristics observer, however, the relationship between reported confidence and orientation estimation performance appears less clear-cut (Figure 2f). This is because of two opposing effects on confidence. When the sample variance across stimulus elements (or external noise) is larger, the perceived amount of noise tends to be higher as well, resulting in lower levels of confidence. Higher levels of internal noise, however, drive confidence levels in the other direction, resulting in reduced visibility and higher levels of confidence. Together, these opposing effects create an inconsistent relationship between confidence and behavioral variability when analyzed within levels of external noise. This also illustrates why studying confidence across stimulus orientations (Figure 4b) may be better suited for the question at hand: because the data are sorted with respect to orientation (which is directly linked to internal noise), the results are easier to interpret.

Our observer models assume that external and internal noise sources are independent. In some cases, internal noise in perceptual computations can have a multiplicative effect on external noise in the input (Dosher & Lu, 1998, 2000), so that the effect of internal noise on behavior scales with external noise variance. Does this impact our findings? Additional simulations showed that when the models include such a scaling effect, this does not qualitatively change the pattern of predictions (Fig. S4).

In summary, an increase in the amount of external noise should result in a decrease in reported confidence, regardless of the strategy to confidence employed by the observer. Predictions diverge between the two models, however, when it comes to changes in internal noise. For an observer who bases confidence on the degree of uncertainty in sensory data, confidence should be inversely linked to variability in behavioral orientation estimates, regardless of the source of noise that creates uncertainty. For an observer who uses heuristic strategies to confidence, however, confidence should paradoxically increase with increasing levels of internal noise. With these predictions in hand, we now turn to the data of human observers to see if their behavior matches that of one of the model observers.

### Human observers

#### Reported confidence

What strategy do human observers employ when reporting confidence? To address this question, human participants viewed a stimulus that consisted of an array of 36 Gabor patches, and reported both the mean orientation of the array, and their confidence in this estimate (Figure 1). Patches were variable in orientation, and five noise levels were used to parametrically manipulate stimulus uncertainty (Experiment 1). In a second experiment (Experiment 2), participants viewed stimuli of a constant noise level with mean orientation ranging between 1° and 180°. Observers had ample practice (at least 350 trials) with the task before the start of either experiment.

Our first analyses focused on the link between external noise, behavioral variability, and confidence (Experiment 1; see *Methods*). As expected, the participants’ orientation estimates were strongly affected by the level of external noise in the stimulus. Specifically, higher levels of orientation noise in the stimulus led to significantly greater variability in reports of perceived orientation (*r* = 0.91, *t*(39 = 14.4, *p* < 0.001) (Figure 3a). Corroborating the predictions of both models, confidence was also highly affected by changes in external noise (Figure 3b). Subjects consistently reported lower levels of confidence with increasing amounts of external noise in the stimulus (*r* = -0.93, *t*(39) = -17.1, *p* < 0.001). Indeed, a direct comparison between confidence and behavioral performance revealed a significant inverse relationship between reported confidence and behavioral variability (*r* = -0.85, *t*(39) = -10.3, *p* < 0.001). Altogether, this indicates that participants were able to meaningfully estimate their own level of confidence in the task.

We then turned to changes in confidence caused by fluctuations in the amount of internal noise. We first focused on sources of noise that are independent of the presented stimulus orientation. For each observer and for each level of external noise, trials were sorted into three bins, based on the level of confidence reported by the observer. Prior to binning, any effects on reported confidence due to physical stimulus properties were removed via linear regression (see *Methods*, Equation 6), so that the bins more closely reflected internal fluctuations in confidence. Behavioral variability and mean level of confidence across all trials in each bin were computed and compared. We considered three possible outcomes. First, our simulations indicated that if observers compute confidence based on the uncertainty in their sensory evidence, as suggested by the Bayesian model, then greater levels of confidence should be linked to more precise (less variable) behavior (Figure 2a, c, e). Second, if observers use a heuristic strategy to confidence based on the perceived amount of noise, then the relationship between confidence and behavioral variability should be inconsistent within external noise levels, as internal and external noise push this relationship in opposite directions (Figure 2b, d, f). Third, if observers use neither of these strategies, then we should observe no reliable link between confidence and behavioral variability at all. Corroborating the Bayesian observer model, we found a positive relationship between confidence and precision in behavior (Figure 3b). Specifically, even within levels of external noise, behavioral orientation estimates were reliably less variable when reported confidence was higher (main effect of confidence, (*F*(2, 18) = 45.1, *p* < 0.001). Additional analyses revealed that higher levels of confidence consistently predicted better behavioral performance when comparing sequentially between confidence bins (high versus medium: *F*(1, 9) = 29.9, *p* < 0.001; medium versus low: *F*(1, 9) = 35.3, *p* = < 0.001). We also found a positive relationship between confidence and behavioral precision for even the lowest level of external noise (i.e. σ_*e*_ = 0.5), when trial-by-trial fluctuations in behavior are likely dominated by internal sources of variance (*r* = 0.51, *t*(19) = -2.6, *p* = 0.016). Control analyses verified that these results do not strongly depend on the number of bins analyzed (main effect of confidence, 2 bins, *F*(1, 9) = 42.2, *p* < 0.001; 4 bins: *F*(3, 27) = 24.4, *p* < 0.001). Thus, these results appear consistent with the Bayesian notion that confidence is computed from the degree of uncertainty in the observer’s perceptual estimates.

To better understand the relationship between confidence and internal noise, we next examined how confidence varied for stimuli at different orientations (Experiment 2; see Methods). Replicating previous findings, behavioral variability was higher for oblique compared to cardinal orientations (*r* = 0.59, *t*(87) = 6.91, *p* < 0.001). Our simulations indicated that if confidence is computed from the precision in sensory evidence, then reported confidence should follow this oblique effect in orientation judgments, with lower levels of confidence for oblique compared to cardinal orientations. If, on the other hand, confidence is based on the perceived amount of stimulus noise, the opposite pattern should emerge and confidence should be higher for oblique than cardinal orientations. Interestingly, comparing confidence between stimulus orientations revealed that participants were more confident about their orientation estimates for oblique than cardinal stimuli (*r* = 0.34, *t*(87) = 3.38, *p* < 0.001), despite lower levels of performance for these stimuli (Figure 4). These findings run contrary to the predictions of the Bayesian observer model and instead provides support for the Heuristics observer model, wherein confidence is based, in part, on external properties of the stimulus.

#### Perceived noise

Our simulations assumed that noise across stimulus elements should be less visible for oblique compared to cardinal orientations. We next validated this assumption in a separate experiment (Experiment 3; see Methods), and used the empirical internal noise values obtained via this experiment to quantitatively predict human confidence reports across orientation stimuli. Human observers were presented with two orientation stimuli in rapid succession. Each stimulus consisted of an array of orientation patches drawn from a Gaussian distribution. The two distributions had the same mean, but differed in their variance, and we used an adaptive staircase to manipulate this variance. Participants were asked to report which of the two stimuli contained the higher level of orientation noise. Repeating the procedure for different mean orientations (from 0 to 165° in 15° steps), and applying an analysis approach based on signal detection theory (Morgan et al., 2008), enabled us to determine the extent to which external noise perception and internal noise values vary across orientation stimuli.

Does human noise perception change across stimulus orientations? If the amount of internal noise is higher for oblique rather than cardinal orientations, then it should be more difficult to detect a change in variance across oblique stimulus elements. Alternatively, if internal noise values remain the same, regardless of orientation, then variance discrimination performance for these orientations should remain the same, as well. Interestingly, our results indicate that performance was better for cardinal than oblique orientation stimuli. Using the approach of Morgan et al. (Morgan et al., 2008), we estimated the amount of internal noise associated with the orientation stimuli (see *Methods*), and found that internal noise was reliably higher for oblique compared to cardinal orientations (Figure 5a; *r* = 0.48, *t(*87) = 5.12, *p* < 0.001). Altogether, this indicates that observers better perceive external stimulus noise around cardinal than oblique orientations – much like we assumed in our modeling work.

To further quantify the Heuristics model’s predictions, we subsequently used the empirical internal noise values as obtained in this experiment to predict human confidence estimates across orientation. The results are shown in Figure 5b. As can be seen from the figure, the human confidence estimates of Experiment 2 are qualitatively well-matched by the model’s predictions for confidence, providing further evidence to suggest that human observers use a heuristic strategy to confidence based on the perceived amount of noise in the stimulus.

## Discussion

Is reported confidence strictly based on a statistical (Bayesian) notion of information, or do observers employ heuristics to estimate their level of confidence in a decision? In the current study, we developed computational models of either strategy, which led to several quantitative predictions that we subsequently tested in behavioral experiments. Our results suggest that observers use a combination of strategies. Specifically, corroborating the Bayesian hypothesis, confidence reliably predicted variability in behavioral orientation judgments when the physical stimulus orientation was held constant. However, when comparing between orientation stimuli, reported confidence was more consistent with the predictions of the Heuristics model, and appeared to reflect the perceived amount of noise in the stimulus. Together, these results suggest that confidence is not always computed from the degree of imprecision in perceptual information but can, instead, also be based on perceptual cues that function as simple heuristics to confidence.

Most work to date on the Bayesian confidence hypothesis has varied physical stimulus properties, such as contrast or duration, to manipulate evidence reliability. Because human confidence ratings often vary in relationship to this manipulation, it was suggested that confidence is computed from the degree of imprecision in perceptual evidence (Barthelme & Mamassian, 2010; Sanders et al., 2016). Our results, however, suggest a more nuanced interpretation: it appears that confidence is not always computed from an explicit representation of perceptual uncertainty, but rather can also be based on visual cues in the stimulus that serve as a proxy to uncertainty. This highlights the importance of studying confidence ratings in the absence of external uncertainty manipulations; for instance, by relying on fluctuations in internal noise that make sensory evidence more or less reliable to the observer (Honig, Ma, & Fougnie, 2020; van Bergen & Jehee, 2019; van Bergen et al., 2015; Walker, Cotton, Ma, & Tolias, 2020).

Why did we manipulate stimulus variance rather than contrast to test our hypothesis? We believe that a contrast manipulation would make it slightly less straightforward to adjudicate between the two strategies. To see why, consider that there is no a-priori reason to assume that estimates of contrast should be more biased for oblique than cardinal orientations (unless one assumes a prior on contrast). In other words, while higher levels of internal noise would create more noisy estimates of contrast for oblique compared to cardinal orientations, the across-trial mean of the contrast estimates would be (roughly) the same, regardless of orientation, and so would reported confidence for the Heuristics observer. For perceived variance, on the other hand, heuristic confidence paradoxically increases for oblique compared to cardinal orientations (Figure 4). Thus, for variance, the predictions more strongly deviate between the Heuristics and Bayesian observers, and this makes it somewhat simpler to test and compare between the two strategies in experiments.

Previous studies have shown that certain physical image properties, such as stimulus contrast or the variability in stimulus elements, can impact confidence judgments (Boldt et al., 2017; de Gardelle & Mamassian, 2015; Herce Castanon et al., 2019; Spence et al., 2016; Zylberberg et al., 2014). Others have suggested that observers use non-normative strategies to estimate confidence (e.g., Adler & Ma, 2018; Filevich, Koß, & Faivre, 2020; Maniscalco & Lau, 2012; Maniscalco, Peters, & Lau, 2016; Peters et al., 2017; Winter & Peters, 2021; Zylberberg et al., 2014), much like we do here. Our work differs from many of these previous studies in that we provide both a Bayesian model and an alternative that explains deviations from optimality. Our normative model explicitly describes the decision-making process and assumes that not only the sensory measurement, but also the level of internal noise varies from trial to trial. This contrasts our approach with other studies that either provided no models at all (Koizumi, Maniscalco, & Lau, 2015; Maniscalco & Lau, 2015; Samaha et al., 2016), used descriptive models (e.g., drift diffusion model, Zylberberg, Fetsch, & Shadlen, 2016), or assumed a fixed level of noise (Maniscalco et al., 2016; Rahnev, Lau, & de Lange, 2011; Rahnev, Maniscalco, Luber, Lau, & Lisanby, 2012). Notable exceptions are several papers that formally model both the Bayesian solution and deviations from optimality (e.g., Zylberberg et al., 2014; Denison et al., 2018; Adler & Ma, 2018; Aitchison et al., 2015; Peters et al., 2017; Winter & Peters, 2021). Some of these previous papers postulated that observers incorrectly estimate the across-trial variance of the stimulus; in other words, observers use uncertainty, but their estimate of the amount of uncertainty is incorrect (see also the discussion in the next paragraph). This is different from our assumption here that, rather than uncertainty, observers base confidence on a property of the stimulus, namely perceived noise, which might itself be estimated correctly. Further, and contrary to previous work, we used a continuous estimation rather than n-alternative forced choice (n-AFC) task. This distinction is important, because the optimal confidence estimate in n-AFC relies on both the sensory measurement and uncertainty, and the effect of the two can be difficult to distinguish in behavior (see also Denison, Adler, Carrasco, & Ma, 2018). For a continuous estimation task, in contrast, confidence should depend on uncertainty only, which makes this task particularly well-suited for the question at hand. In addition, and perhaps as a consequence of the n-AFC task, most work to date has focused exclusively on heuristic explanations in terms of the sensory measurement (e.g., “positive evidence” heuristic in Maniscalco et al., 2016, a “quadratic” heuristic in Adler & Ma, 2018, or a “inflexible criterion” or “variance misperception” suboptimality in Zylberberg et al., 2014), with little focus on uncertainty per se, whereas our Heuristics model directly addresses noise or uncertainty perception. Note that this difference between n-AFC and continuous estimation also makes a common measure of metacognitive sensitivity, meta-d’ (Maniscalco & Lau, 2012), inapplicable in our study, as this metric was developed explicitly for the n-AFC task.

Many previous studies have suggested that observers in perceptual tasks are often overconfident; that is, their confidence levels are too high compared to their objective performance (e.g., Baranski & Petrusic, 1999; Koriat, 2011; Pleskac & Busemeyer, 2010; Winter & Peters, 2021; see Rahnev & Denison, 2018, for a recent review). Zylberberg et al. (2014) suggested that such biases could partly stem from an underestimation of the across-trial variance of noisy stimuli. Similarly, Peters et al. (2017) proposed that observers have higher internal uncertainty due to TMS stimulation, but do not take all of this uncertainty into account, resulting in an overestimate of confidence. Thus, in both studies, observers are thought to appropriately rely on their internal uncertainty, but to incorrectly estimate its value. Our work, on the other hand, indicates that observers do not underestimate uncertainty *per se*, but rather appear to use the amount of perceived noise as a heuristic to confidence. As such, our findings point to an important alternative explanation for why observers sometimes tend to be ‘overconfident’ in their judgments.

Experimental evidence (e.g., Girshick, Landy & Simoncelli, 2011) suggests that human observers apply in many perceptual decisions a prior distribution that favors cardinal over oblique orientations, matching the orientation statistics of the natural environment. Would this impact our conclusions? Note that if such a prior were a dominant influence on our participants’ confidence reports, we would have expected their confidence to increase near cardinal orientations (where the natural orientation prior is more concentrated), and decrease around the obliques. Instead, we find the exact opposite pattern in confidence as a function of orientation. This suggests that a non-Bayesian (or anti-Bayesian) strategy, like our Heuristics observer model, must be the dominant effect here. If we supposed that a naturalistic prior nevertheless had some influence on our participants’ confidence reports, then an even stronger anti-Bayesian effect would be required in order to explain our findings.

An implication of our findings is that neural populations encoding for confidence should receive information about both perceived noise and sensory uncertainty. Previous work from our lab has shown that sensory uncertainty can be decoded from activity in early visual areas – even in the absence of explicit uncertainty manipulations (van Bergen & Jehee, 2018, 2019; van Bergen et al., 2015), and we predict that this decoded uncertainty should also be reflected in higher-level areas, such as the anterior cingulate cortex and prefrontal cortex, that have been implicated in confidence (Bang & Fleming, 2018; De Martino, Fleming, Garrett, & Dolan, 2013). It will be interesting for future studies to test this prediction.

While perceptual decisions often conform to the principles of Bayesian theory (reviewed in e.g., Knill & Pouget, 2004), in economic and cognitive decision-making heuristic strategies are common currency: numerous cognitive and behavioral economic studies have shown that human observers take heuristic shortcuts to decision confidence (see Lichtenstein et al., 1977). Our results, together with other findings (e.g., Aitchison et al., 2015; Zylberberg et al., 2014), suggest that confidence about perceptual decisions may in some instances share these aspects with higher-level decision-making. It will be interesting for future studies to further investigate the extent to which perceptual and cognitive strategies in human decision-masking converge.

The Heuristics observer model bases confidence on the perceived amount of external orientation variability in the stimulus. In order to arrive at this estimate, the observer accounts for the fact that some of the variability in sensory measurements can be ascribed to internal sources of noise. When computing confidence, however, this observer ignores internal noise variance and bases confidence on the perceived amount of external noise alone. This is patently suboptimal, and is the reason for this observer’s anti-Bayesian confidence patterns. Why would an observer incorporate internal noise variance in one case, and ignore it in another? While answering this question is beyond the scope of the present study, one intriguing possibility is that the estimation of perceived noise is a strictly perceptual computation, the outcome of which is subsequently sent to higher-level areas involved in the computation of confidence. Neurons involved in low-level perception may have computational access to the internal noise variance associated with their sensory inputs, while such detailed information is not always communicated towards areas involved in metacognition or decision-making. This would be consistent with the aforementioned view that perceptual confidence may have strong parallels with confidence in higher-level decisions. However, it is also worth noting that the heuristics observer is only intended to model the specific deviation from optimality that we observed here. It is not intended as a comprehensive model of human metacognition, which may well incorporate knowledge of internal noise in confidence estimates – perhaps especially when “cheap” heuristics based on external stimulus statistics are not available. This interplay between heuristics and more ‘Bayes-aligned’ computations will be an interesting focus for future research.

In conclusion, the current results suggest that observers use a dual strategy to decision confidence: not only do they utilize an explicit representation of sensory uncertainty (as postulated by normative theories), but they also consider various stimulus properties as simple cues to confidence. This presents an important word of caution to normative studies of decision making, and highlights the need for studying confidence in the absence of explicit uncertainty manipulations.

## Acknowledgments

This work was supported by European Union Program FP7-PEOPLE-2013-ITN Grant 604063-IDP (A.B.), and by European Research Council Starting Grant 677601 (to J.F.M.J.).

## Citation gender diversity statement

The gender balance of papers cited within this work was quantified by manual gender determination for the first and the last authors. Among the 52 cited works that had named

authors, there were 42 unique first authors (19% woman) and 32 unique last author (22% woman). This is below the base rate (approximately 30% among faculty members) observed in the field based on the data from Vision Science Society meetings (Cooper & Radonjic, 2016).

## Supporting Materials

### Noise detection model

The third model observer uses the same decision-making process as the Bayesian and perceived-noise model to report the orientation of the stimulus. However, rather than calculating confidence from uncertainty or the perceived amount of noise, this model assumes that observers base their confidence on the degree to which they detect noise in the stimulus. More specifically, this model observer first computes the sample variance across the measured stimulus elements and then calculates the probability that external noise was present in the stimulus. Confidence is based on this measure of noise visibility.

Because the measurements of individual stimulus elements are drawn from a Gaussian distribution 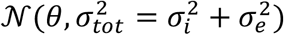, their sample variance is distributed as a scaled χ^2^ distribution with *k*-1 degrees of freedom (where *k* is the number of measurements on a single trial):

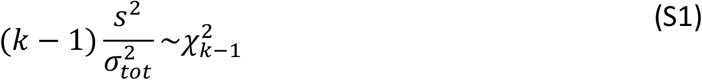

We assume that for each trial, the observer computes the sample variance *s*^2^ across the measured stimulus elements (i.e.,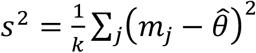) and then uses this to calculate the probability that the stimulus contained no external noise:

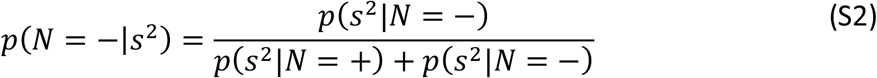

The probability of observing sample variance *s*^2^ when the stimulus contains external noise (i.e., *p*(*s*^2^|*N* = +)) can be computed by integrating over all possible external noise values:

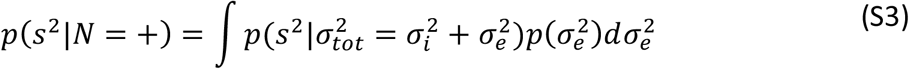

When the stimulus contains no external noise, the probability of observing *s*^2^ is simply:

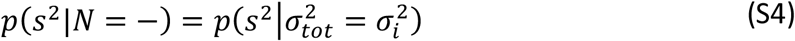

For the noise-detection model observer, confidence on a given trial is directly related to the probability that there is no external noise:

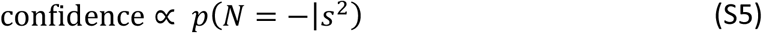

### Simulations

In our analyses, we assumed that 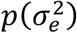 was equivalent to the distribution of discrete noise levels used in the experiments. This turns the continuous integral in Equation S3 into a discrete sum, which was computed numerically. We then simulated the sample variance the same way as described in the Methods section of the main text and computed trial-by-trial confidence estimates using Equation S5. See Figures S1 and S2 for the model’s predictions.

**Figure S1.**
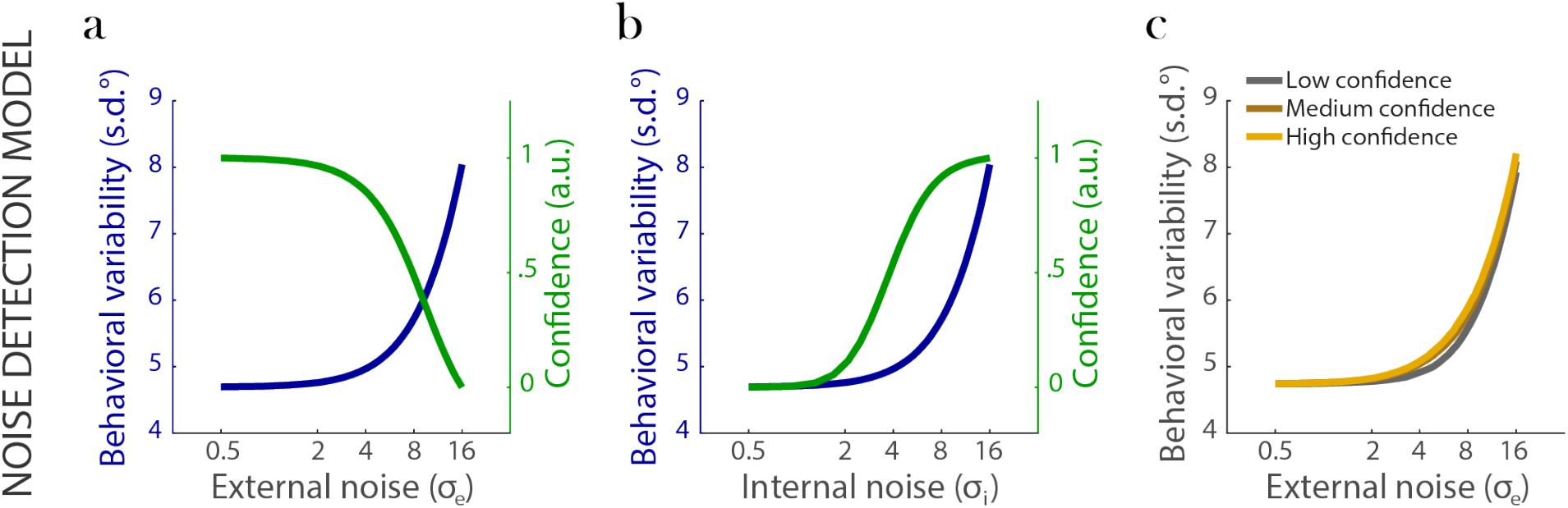
Predictions of the noise-detection model observer. Behavioral variability (blue) increases with increasing levels of external **(a)** and internal **(b)** noise. Confidence (green) nicely tracks the decrease in behavioral precision that occurs with increasing levels of external noise **(a)**. However, for the Heuristics observer model that bases confidence on the ability to detect external noise, **(b)** confidence paradoxically increases with larger amounts of internal noise. **(c)** Behavioral variability for three levels of confidence. For each simulated level of external noise, trials were sorted into three bins of increasing confidence, and behavioral variability was computed across all trials in each bin. The effects of external and internal noise go in opposite directions, resulting in an inconsistent relationship between confidence and behavioral precision. In all figures, the x-axis is plotted using a logarithmic scale.

**Figure S2.**
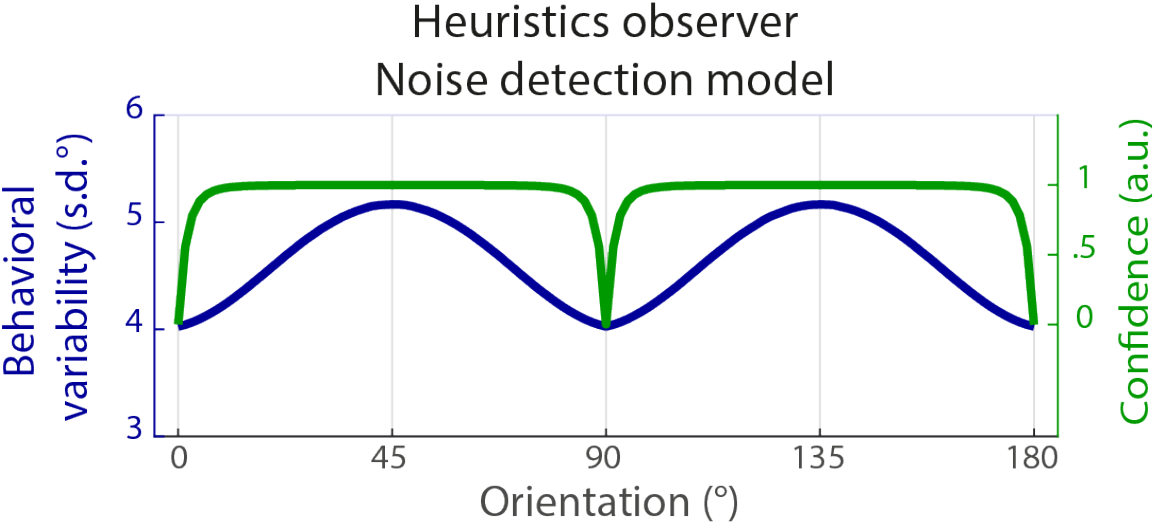
Behavioral variability (blue) and reported confidence (green) for the Heuristics noise-detection model observer. Confidence predicts *poorer* behavioral performance: when behavioral variability increases due to higher levels of internal noise (oblique orientations), confidence is higher.

**Figure S3.**
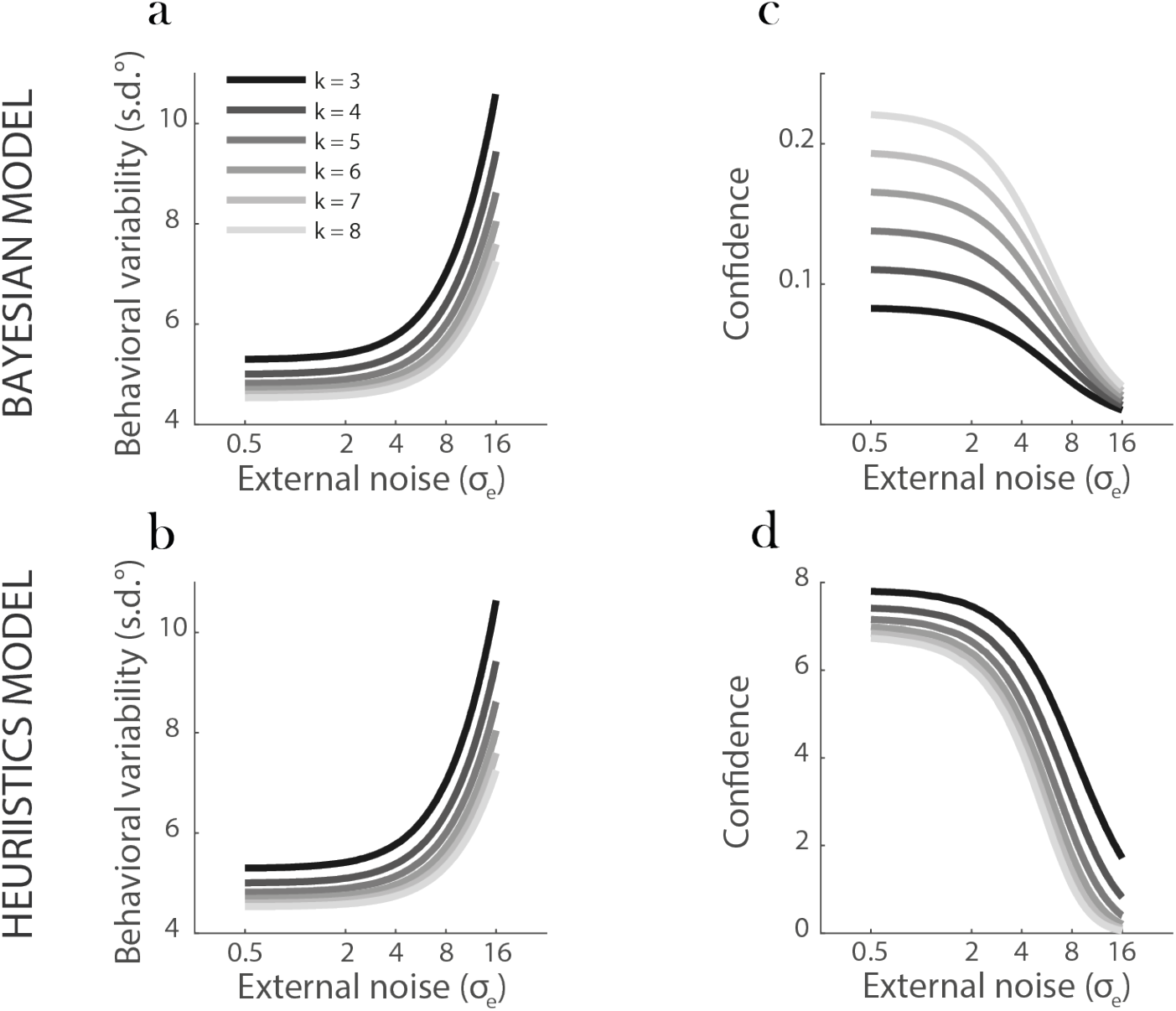
Predictions for the Bayesian and Heuristics model observers for various values of *k* corresponding to the number of patches sampled by the observer. The parameter *k* merely shifts (or scales) the behavioral-variability and confidence curves up or down, but does not change the qualitative pattern of results (predictions) of either observer model. (**a-b**) Behavioral variability as a function of external noise and *k* for the Bayesian (**a**) and Heuristics (**b**) model observer. For both observers, behavioral variability increases with increasing levels of external noise under all values of *k*. (**c-d**) Confidence as a function of external noise for various levels of *k* for either the Bayesian (**c**) or the Heuristics (**d**) model observer. Confidence decreases with increasing levels of external noise and *k* only shifts the amplitude of the predictions. Behavioral variability and confidence were computed using the same procedure as in Figure 2a-d. In all figures, the x-axis is plotted using a logarithmic scale.

**Figure S4.**
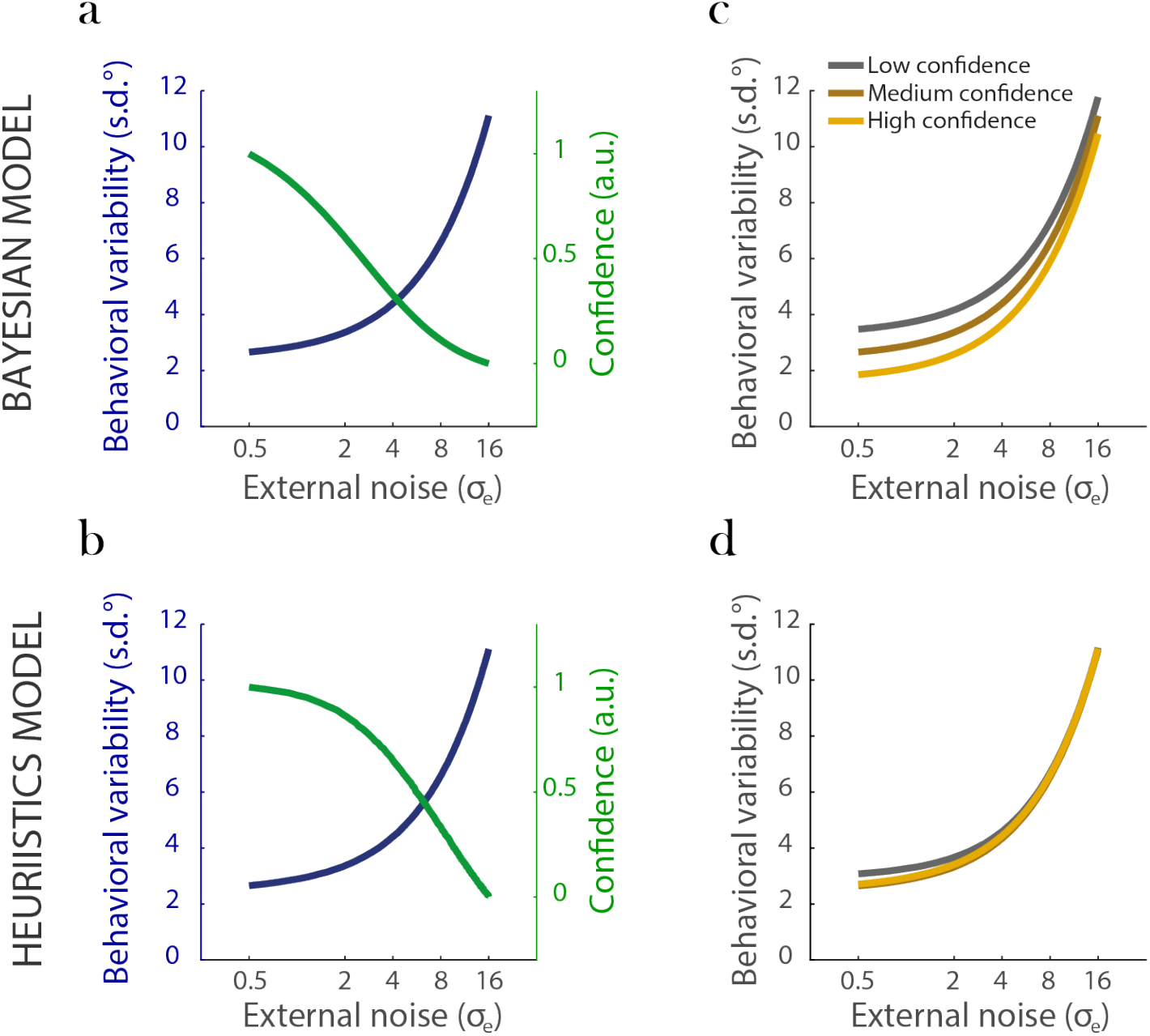
Predictions for the Bayesian (**a-b**) and Heuristics (**c-d**) model observer where internal noise scales with the overall variance of the stimulus, rather than being constant across external noise levels (cf. Figure 2). Internal noise was defined as 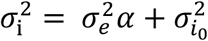, where σ was set to 120 linearly spaced values from 0.5° to 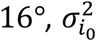 was set to 6° (**a** & **b**), or either low, medium or high (i.e., 4°, 6° and 8°, **c** & **d**), and *α* = 1. Other details are as described in *‘Simulations’*. Considering a level of internal noise that scales with external noise does not strongly affect the qualitative predictions of either model observer. That is, replicating Figure 2, (**a-b)** Behavioral variability increases as a function of external noise (blue lines) and confidence well predicts these changes in behavioral performance (green lines). (**c-d**) Predictions when trials are sorted with respect to reported confidence. For each level of external noise, trials were sorted into three bins of increasing confidence, and behavioral variability was computed across all trials in each bin. The level of confidence reported by the Bayesian observer (**c**) correctly predicts behavioral variability: when the observer reports high confidence, orientation estimates tend to be less variable. For the Heuristics observer (**d**), there is no fixed relationship between confidence and behavioral performance, as the effects of external and internal noise compete. In all figures, the x-axis is plotted using a logarithmic scale. We similarly verified that the qualitative pattern of predictions did not strongly depend on the value of *α* (where we used *α* ∈ {0.2, 0.4, 0.6, 0.8, 1.0}; data not shown).

